# Full-length COI barcodes improve eDNA metabarcoding data denoising relative to mini-barcodes

**DOI:** 10.64898/2026.07.03.736260

**Authors:** Marius H Eisele, Sirle Varusk, Kaarel Sammet, Ali Hakimzadeh, Martin Metsoja, Leho Tedersoo, Khairiah Mubarak Alwutayd, Paula Arribas, Carmelo Andújar, Brent C Emerson, Sten Anslan

## Abstract

Animal COI (mitochondrial cytochrome oxidase I) metabarcoding of environmental DNA (eDNA) is increasingly used to assess biodiversity in complex substrates such as soil. However, due to read-length constraints of second-generation sequencing platforms, mini-barcodes have been used instead of the full barcode region. Long-read sequencing technologies now enable the recovery of full-length barcode sequences, and are more commonly applied for studying microbes, but their use for metabarcoding the full-length standard COI barcoding region in animals remains limited. In this study, we compared three COI amplicon sets – 313 bp, 660 bp, and 1,256 bp – amplified from soil eDNA samples and sequenced using Illumina and PacBio platforms to evaluate their overall concurrence, the effectiveness of identifying nuclear mitochondrial DNA segments (NUMTs) and chimeras, as well as their respective taxonomic resolution. The long-read datasets exhibited a higher identification rate of NUMTs and true chimeras, suggesting that longer sequences improve the detection of noise in COI metabarcoding data, thereby reducing the occurrence of spurious taxa. Taxonomy assignment confidence was similar between the 313 bp and 660 bp datasets, whereas extending the amplicon beyond the standard COI barcode region (1,256 bp) reduced confidence, likely because longer reads extend into regions poorly represented in barcode reference databases. Despite substantially lower sequencing depth in the 660 bp dataset, per-sample OTU richness did not differ significantly from that recovered with the Illumina 313 bp amplicon set. Similarly, the relationships between samples were strongly correlated across the detected OTU communities, indicating consistent ecological interpretations between short and long amplicons. We conclude that the standard ∼658 bp COI barcode is an optimal marker for soil animal metabarcoding from eDNA, balancing target recovery, artifact detection, taxonomic assignment and ecological interpretability. As COI eDNA metabarcoding becomes increasingly used in biodiversity assessment and is increasingly adopted in large-scale monitoring initiatives, this study provides methodological guidance for improving the robustness of soil animal community biomonitoring.

## Introduction

Metabarcoding has emerged as a powerful and cost-effective tool for rapid characterization of organisms from environmental or bulk samples through DNA sequencing of standardized genetic amplicons. It has been extensively applied to study the community compositions across the tree of life - from microbial ecology and plant biology to animal and marine studies - offering a unifying framework for comparative biodiversity analysis (e.g., Compson *et al*. 2020, Cowgill *et al*. 2025, Deiner *et al*. 2017, Taberlet *et al*. 2012). The expanding scope and complexity of metabarcoding studies have been made possible by continuous advancements in sequencing technologies, which have steadily improved throughput, accuracy, and cost-efficiency (Bush *et al*. 2019). Since the advent of high-throughput sequencing-by-synthesis technology by Illumina, this method of sequencing has become the dominant approach in the field of DNA metabarcoding (Uhlen & Quake 2023). It offers a viable solution for ultra-high sequencing depth but is currently (as in early 2026; but see Illumina, 2025) restricted to produce a maximum of 2 × 300 base pair (bp) sequences with the Illumina MiSeq. Although 2 × 500 bp sequencing is technically emerging, it is not yet widely offered and is currently approximately 1.5 times more expensive than 2 × 300 sequencing (Genomics and Cell Characterization Core Facility [GC3F], 2026). For animals, the standard DNA barcoding region is a ∼658 bp fragment of the mitochondrial cytochrome c oxidase subunit I (COI or cox1) gene (Hebert *et al*. 2003), which exceeds the sequencing length capabilities of widely available second-generation sequencing technologies (including Illumina). Therefore, metabarcoding studies have shifted to so-called mini-barcodes, which represent a nested subset of a standard barcoding region to meet the sequencing requirements of the second-generation sequencing instruments.

More recently, third-generation real-time single-molecule sequencing machines, which emerged about a decade ago, have enabled the sequencing of much longer DNA molecules, reaching tens or even hundreds of thousands of base pairs (Jain *et al*. 2018, Rhoads & Au 2015). Generally, there is a trade-off between sequence length and sequence quality (Amarasinghe *et al*. 2020). However, Pacific Biosciences’ (PacBio) third-generation sequencing has overcome this limitation through multiple passes of circular consensus sequencing, which can produce up to 25 kb high-quality (HiFi) reads (Tedersoo *et al*. 2022; Wang *et al*. 2025). This has allowed a return to utilizing full-length barcodes in metabarcoding approaches, which is being embraced especially in studies targeting microbes, such as bacteria (Buetas *et al*. 2024; Gao *et al*. 2024) and fungi (Kolarikova *et al*. 2021; Mikryukov *et al*. 2023; Tedersoo & Anslan 2019), but also other microeukaryotes (Jamy *et al*. 2020; Latz *et al*. 2022; Mikryukov *et al*. 2026). For taxa in the animal kingdom, third-generation sequencing has been primarily applied to scale up individualized-specimen barcoding efforts (Cuber *et al*. 2023; Hebert *et al*. 2025; Srivathsan *et al*. 2024), but has only been used to a limited extent in metabarcoding studies (e.g., Lin *et al*. 2025; Varusk *et al*. 2025).

When amplifying mitochondrial DNA amplicons such as COI, another important consideration in a metabarcoding dataset is the presence of nuclear mitochondrial DNA segments (NUMTs). NUMTs are fragments of mitochondrial DNA that have been transferred to the nuclear genome and are frequently co-amplified during PCR when using universal mitochondrial primers (Bensasson et al. 2001). Following nuclear integration, NUMTs may accumulate mutations at a rate decoupled from genuine mitochondrial evolution (Bensasson et al. 2001; Song et al. 2008), meaning they risk inflating diversity estimates or producing erroneous taxonomic assignments if not identified and excluded. Artefactual recombinants generated during PCR - chimeric sequences - pose a similar problem. If not filtered out, they introduce spurious haplotypes that inflate diversity and taxonomic inferences (Zinger et al. 2019; Edgar et al. 2011). Removal success of both artefact types are, to some extent, shaped by the amplicon length. Short reads may lack sufficient sequence information to discriminate genuine mitochondrial sequences from NUMTs or to resolve chimeric breakpoints, given the reduced number of informative sites available across the fragment (Edgar et al. 2011). Longer amplicons, by spanning a greater proportion of the gene, are more likely to expose NUMT-associated frame disruptions, internal stop codons, or the conflicting phylogenetic signals characteristic of chimeric recombination (Porter & Hajibabaei 2021; Hakimzadeh et al. 2025). The key trade-off, however, is that longer amplicons may also increase the probability of incomplete extension during PCR, which can itself promote chimera formation, particularly when amplification efficiency varies across templates (Edgar et al. 2011, Tedersoo et al. 2021). Despite this trade-off, long-read sequencing approaches may offer meaningful advantages in detecting and filtering both artefact types during downstream bioinformatic processing.

Given the growing use of COI-based animal eDNA metabarcoding and the limited number of studies directly comparing alternative sequencing strategies, there is a need for empirical assessments that can inform methodological choices and contribute to the development of more standardized workflows. This study examines the overall concordance between short-and long-read sequencing datasets of COI amplicons generated from soil environmental DNA (eDNA), as well as their effectiveness in identifying NUMTs and chimeras, and taxonomic resolution. By evaluating commonly used short-read and emerging long-read approaches, we assess the advantages, limitations and implications of each strategy for biodiversity inference from soil eDNA. Amplicon libraries for the short-read sequencing (Illumina) were generated with commonly used primers for short-read metabarcoding of fauna targeting a 313 bp fragment in the COI gene, whereas the amplicon libraries for long-read sequencing (PacBio) were generated using standard barcoding primers targeting ∼658 bp (Folmer region) and a newly designed primer targeting a 1,256 bp of the COI gene.

## Materials and Methods

### Sampling

Soil samples used in this study are a subset of samples from Anslan *et al*. (2021). Accession numbers from the Sequence Read Archive (SRA) are listed in **Table S1**. Briefly, soil samples were collected from various terrestrial ecosystems – shrublands (former agricultural land, or previously managed grassland), young forests (stand age <20 years) and forests. Nine soil cores (⌀ of 5 cm and 10 cm deep) from each 3 m × 3 m plot were collected, approximately 15 g of soil was scraped from the core sides and pooled in a Zip-Lock plastic bag. The composite sample was thoroughly mixed and stored at-80 °C.

### Molecular analyses

Detailed molecular analyses are outlined in Anslan *et al*. (2021). In short, frozen soil samples were crushed and dried in a drying cabinet at 35 °C for 24 h. The dried samples were thoroughly homogenized, and a 0.25 g subset was used for DNA extraction. DNA extraction was carried out with a Thermo Scientific KingFisher Flex robot and the MagAttract PowerSoil kit (Qiagen Inc., Hilden, Germany), following the manufacturer’s guidelines. DNA extraction also included a blank extraction (without soil).

Polymerase chain reactions (PCRs) were carried out using three sets of primers to generate amplicon libraries targeting 313 bp, 660 bp and 1,256 bp of the mitochondrial cytochrome oxidase I (COI) gene region (**Figure 1**). PCRs for the 313 bp amplicons were performed with primers mlCOIintF (5′ GGW ACW GGW TGA ACW GTW TAY CCY CC) Leray *et al*. (2013) and jgHCO2198 (3′ TAI ACY TCI GGR TGI CCR AAR AAY CA) (Geller *et al*. 2013). This primer combination is commonly used in short-read metabarcoding studies (Elbrecht *et al*. 2019, Wangensteen et al. 2018). PCRs for the 660 bp amplicons were performed with primers LCO1490 (5′ GGT CAA CAA ATC ATA AAG ATA TTG G) and HCO2198 (3′ TAA ACT TCA GGG TGA CCA AAA AAT). This set of primers represents commonly used barcoding primers in high-throughput sequencing workflows (Liu et al. 2017), but has also been demonstrated to be useful in metabarcoding of soil fauna directly from soil DNA (Varusk *et al*. 2025). PCRs for the 1,256 bp amplicons were performed with primers jgLCO1490m (5′ TCI ACI AAY CAY AAR GAY ATT G) and CM1300R (3′ GAR TAW CGT CGN GGT ATN CC). Primer jgLCO1490m is a shortened version of jgLCO1490 (Geller *et al*. 2013) to optimize PCR compatibility with the newly designed primer CM1300R. This new primer was based on Collembola and Acari COI regions extracted from EMBL v143 (Kanz et al. 2005) and available GenBank mitogenomes (Sayers et al. 2019). All primers (except jgHCO2198) included unique indexes (8-12 bp) per-sample for multiplexing. The PCR mixture (total volume of 25 µl) consisted of 5 µl of 5× HOT FIREPol Blend Master Mix (Solis Biodyne in Tartu, Estonia), 0.5 µl of each forward and reverse primer at a concentration of 20 mM, 1 µl of DNA extract, and 18 µl of ddH_2_O. PCR cycling conditions for generating amplicon library targeting 313 bp of the COI region included an initial denaturation at 95 °C for 15 min (hot-start for HOT FIREPol® Blend Master Mix); 35 cycles of denaturation for 30 s at 95 °C, annealing for 30 s at 57 °C, elongation for 1 min at 72 °C; and final extension at 72 °C for 10 min and storage at 4 °C. PCR conditions for primer pair LCO1490-HCO2198 (targeting 660 bp) were as follows: initial denaturation at 95 °C for 15 min; 5 cycles of 95 °C for 30 s, 45 °C for 1 min, and 72 °C for 1 min; followed by 35 cycles of 95 °C for 30 s, 51 °C for 1 min, and 72 °C for 1 min, with a final extension at 72 °C for 10 min. Latter PCR conditions were also used for primer pair jgLCO1490m-CM1300R with the adjustments of 8 cycles for 45 °C and 32 cycles for 51 °C annealing step, and extending the elongation time to 1 min 30 seconds.

**Figure 1.**
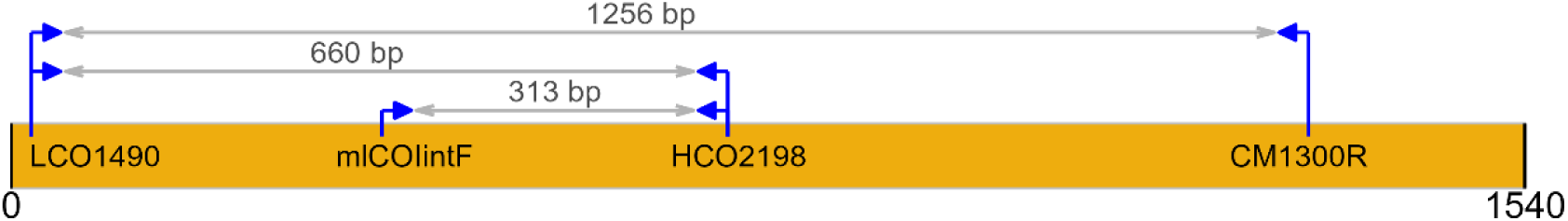
Primer binding sites and amplicon lengths (without primers). Primers mlCOIintF and jgHCO2198 were used to generate 313 bp amplicons, LCO1490 and HCO2198 for 660 bp amplicons, and jgLCO1490m and CM1300R for 1,256 bp amplicons. The primer map is based on *Drosophila yakuba* COI gene (KF824885; Llopart *et al*. 2014).

Two PCR reactions were performed per sample and pooled for PCR product verification via 1% agarose gel electrophoresis. Samples that yielded no product were re-amplified using 2 µl of DNA. Negative PCR controls, consisting of ddH2O without any DNA template, were employed to detect any apparent contamination during sample preparation for PCR. Positive controls (synthetic DNA molecules) were used to assess PCR efficiency. PCR products were normalized based on visual inspection of band strength on a 1% agarose gel and then pooled. We used the following criteria: no band (i.e., negative PCR control and DNA extraction control) = 10 µl; faint band = 7 µl; medium band = 3 µl; strong band = 1 µl. The pooled amplicon library consisting of 313 bp COI fragments was sequenced with Illumina NovaSeq 6000 paired-end sequencing (2 × 250 bp) at Novogen (UK). Amplicon libraries consisting of 660 bp and 1,256 bp COI fragments were sequenced with PacBio Sequel II platform at the Norwegian Sequencing Centre (Norway). Sequencing libraries for respective sequencing platforms (including adapter ligation) were prepared by the service providers.

### Bioinformatics

Raw paired-end Illumina and single-end PacBio data were processed using a pre-defined DADA2 v1.34 (Callahan *et al*. 2016) pipeline as implemented in PipeCraft2 v1.1.0 platform (Anslan *et al*. 2017). This included cutting the primers with cutadapt v4.4.0 (Martin 2011) by allowing a maximum of two mismatches to the primers and a minimum overlap of 19 bp. For 313 and 660 bp amplicon data, sequences exceeding the maximum expected error rate of two and any ambiguous bases (Ns) were discarded during the quality filtering step. The expected error rate is a sum of per-base error probabilities along the sequence. With increasing sequence length, the sum of error probabilities naturally increases because each base has an error probability encoded in its quality score. Therefore, for the 1,256 bp amplicon data, the maximum expected error rate was increased to five. Quality filtered sequences were denoised using the DADA2 denoising algorithm, where the error estimation function for Illumina data was *loessErrfun* and *PacBioErrfun* for PacBio sequences (detect singletons set to true for PacBio; other denoising parameters were kept as default). Putative chimeras were filtered out using the *consensus* method within DADA2. The resulting amplicon sequence variant (ASV) tables were processed with UNCROSS2 (Edgar 2018; within PipeCraft2) to filter out potential tag-jump events (f = 0.03, p = 1). Additionally, minimum and maximum requested sequence lengths for the 313 bp amplicon set was 307 and 319 bp, respectively; for 660 bp amplicons, 654 and 666 bp; and for 1,256 bp amplicons 1,250 and 1,262 bp (± 6 bp for all expected amplicon lengths). ASVs were classified using SINTAX (Edgar 2016) with cutoff of 0.8 in PipeCraft2 (wraps vsearch v2.29.4; Rognes *et al*. 2016) where BOLDistilled (Prosser *et al*. 2025; November 2025 version) served as a reference database (includes 1,742,793 Animalia, 8,250 Protista, 7,941 Bacterial and 831 Fungal sequences). The ASV tables per amplicon set were filtered to remove putative contaminants based on the ASVs in the negative control samples using *decontam* v1.26.0 (Davis et al. 2018) R package (method = prevalence, threshold = 0.1).

### Quantification of authentic and non-authentic OTUs

To further reduce noise in the dataset, we applied metaMATE v0.5.7 (Andújar et al. 2021; https://github.com/tjcreedy/metamate), a tool designed to filter non-authentic sequences such as nuclear mitochondrial pseudogenes (NUMTs). For metaMATE, we included only ASVs classified as Animalia at the bootstrap level >= 0.8 (excluding Chordata). Animalia ASVs were clustered into OTUs in PipeCraft’s “ASV to OTU” module, which performs clustering with vsearch (settings: 97% sequence similarity, --iddeff = 2).

We employed an ‘OTU mode’ of metaMATE where OTUs are classified as verified authentic when at least one ASV of an OTU is an exact match to a reference sequence, and as verified non-authentic when every ASV within an OTU either lay outside the expected length range (allowed base variation, --basesvariation = 6) or included stop codons. Other OTUs receive an ‘unclassified’ category by metaMATE. Because most references sequences in the Barcode of Life Data System (BOLD) database (Ratnasingham & Hebert 2007; November 7, 2025 version) are much shorter than the 1,256 bp amplicon set (thus, the verified authentic OTUs classification is hampered), the 1,256 bp sequences were truncated to the first 660 bp (5′ end) and dereplicated prior to executing metaMATE. The full BOLD database (downloaded in December 2025) was used as the reference dataset to maximize detection of authentic mitochondrial sequences. Sequences shorter than 600 bp were removed from the reference database prior to analysis, keeping around 15 million sequences. Per amplicon set, metaMATE was then executed with the Animalia OTUs datasets using the *--filter-adaptive* mode, which tests a range of per-sample abundance thresholds and evaluates the retention and rejection of authentic and non-authentic OTUs (with authentic and non-authentic proportions extrapolated from the verified fraction to the entire dataset) at each threshold. This allows the selection of a threshold at which a defined proportion of non-authentic OTUs are removed, here set to 95% (*--percentile 0.95, --criteria verified_removed*). Per-sample rarefaction curves of observed OTUs (after metaMATE filtering) indicated that sequencing depth was generally sufficient for the 313 bp and 660 bp amplicon sets, with most samples approaching asymptotes (Figure S1). In contrast, some per-sample rarefaction curves in the 1,256 bp dataset had not yet reached a plateau, indicating that higher sequencing effort would yield additional OTUs for some samples.

To directly compare OTU status assignments by metaMATE (authentic, non-authentic and unclassified) we performed clustering of the ASVs (only Animalia ASVs) from all three datasets. To reduce complexity of the analysis, we first computed (a) the shared OTUs of the 313 bp and 660 bp amplicon sets and then, (b) the shared OTUs between the 660 bp and 1256 bp amplicon sets. Before clustering 313 bp and 660 bp ASVs, the terminal 300 bp region was extracted from each ASV sequence in both datasets using seqkit2 (v2.2.0; Shen *et al*. 2024). For clustering 660 bp and 1,256 bp ASVs, we used the 1,256 bp sequences that were truncated to the first 660 bp (same set as subjected to metaMATE). Clustering of (a) and (b) was performed with vsearch (--cluster_size) at 97% identity (--id 0.97, --iddef 2). We focused on unified OTUs (clusters) containing ASVs from both amplicon sets and assessed agreements in metaMATE-assigned OTU status categories between 313 bp and 660 bp datasets (a). Because the ASVs for clustering (a) were clipped to 300 bp, there were several cases where multiple OTUs formed by ASVs (with the original read lengths) were collapsed into a single unified OTU. This resulted in 902 unified OTUs (899 for 660 bp and 13 for 313 bp) that showed mixed OTU status within a dataset, meaning that the ASVs from either 313 bp or 660 bp dataset did not share a common metaMATE OTU status category (e.g. unclassified and non-authentic co-occurred within the 660 bp subset of a unified OTU). Those unified OTUs with mixed OTU status categories were excluded from the comparison, keeping 1,973 unified OTUs for the following analysis. Additionally, we compared taxonomic assignment confidence between the amplicon sets using only the shared unified OTUs that were represented in the corresponding final OTU sets (after all filtering steps). Unified OTUs from 660 bp and 1,256 bp datasets were only used for comparing the taxonomy assignment confidence.

### Identification of false-positive and false-negative chimeras

Additionally, we explored the chimera filtering efficiency across different amplicon sizes by using the BlasCh module (Hakimzadeh et al. 2025) in PipeCraft v1.2.0. BlasCh applies BLASTn (Camacho et al. 2009) search for the putative chimeric sequences (ASVs filtered out during the DADA2 chimera filtering process) to identify false-positive chimeras (sequences misclassified as chimeras). Here, BOLDistilled served as a reference database. Query ASVs that demonstrated at least 95% sequence identity with at least 95% query coverage against reference sequences were interpreted as likely representing authentic biological sequences that had been incorrectly flagged as chimeric. Here, the false-positive chimeras were not reincluded in the final dataset but were used to compare the relative proportions of false-positive chimeras across different COI amplicon sets. Additionally, BlasCh was applied to the final metaMATE filtered OTUs of each amplicon set for screening for false negative chimeras (potentially chimeric sequences that were not detected by the chimera filtering process). OTU representative sequences with multiple alignments to different target sequences in the reference database were considered as false negative chimeras. Performance was evaluated using the precision (ranging from 0 to 1), calculated as true positive/ (true positive + false positive).

### Statistics and visualization

We used Wilcoxon rank-sum tests (with Bonferroni correction) to test the differences in the per sample OTU richness among the three amplicon sets (313 bp, 660 bp and 1,256 bp). OTU richness per sample was calculated from the OTU data that passed metaMATE filtering (final OTU tables). To assess congruence among community composition ordinations derived from the three COI amplicon sets, we performed Procrustes analysis (9999 permutations) using the vegan package (v2.6.10; Oksanen et al. 2007) in R (v4.4.3; R Core Team 2025). Ordinations were generated using non-metric multidimensional scaling (NMDS) with the metaMDS function in the vegan package. Prior to ordination, OTU tables were presence-absence transformed and pairwise sample dissimilarities were calculated using the Jaccard index with the vegdist function in vegan. The Procrustes correlation coefficient (Procrustes R) was used as a measure of concordance between ordinations, with values closer to 1 indicating greater similarity in the multivariate community structure recovered by each amplicon set.

In addition to these statistical comparisons, we visually summarized patterns in OTU composition and sampling completeness. Pie charts of verified authentic OTU compositions were generated using the matplotlib package (v3.10.8; Barrett et al. 2005) in Python (v3.12.9; Python Software Foundation, 2024), while rarefaction curves were produced with the scikit-bio package (v0.7.2; Aton et al. 2025) and also visualized using matplotlib. Other figures were generated with ggplot2 v4.0.0 (Wickham 2011) in R.

## Results

Across the 34 soil samples, the average raw sequencing depth per sample was 656,990 for the 313 bp amplicon (sequenced with Illumina NovaSeq platform), whereas the average sequencing depths for PacBio datasets were 27,987 reads for the 660 bp amplicon and 24,747 reads for the 1,256 bp amplicon. Following the DADA2 pipeline and length filtering, the remaining average number of sequences per sample was 548,371 (74.4% retained) for 313 bp amplicon dataset, 23,068 (81.9% retained) for the 660 bp dataset, and 7,660 (30.7% retained) for the 1,256 bp dataset (Figure S2). In total, 95,402, 115,124 and 16,653 ASVs were documented in 313 bp, 660 bp and 1,256 bp datasets, respectively (Table 1; Figure S2). From this set of ASVs, the proportion of the target group (Animalia) was 52.04%, 28.88% and 57.02% for 313 bp, 660 bp and 1,256 bp amplicon sets, respectively (Table 1). On average, Animalia ASVs accounted for 18.5%, 74.8% and 17.6% of the raw number of sequences per sample in the 313 bp, 660 bp and 1,256 bp amplicon datasets, respectively (Figure S2). Clustering of Animalia ASVs at a 97% similarity threshold yielded 20,728 OTUs for the 313 bp amplicon set, 22,792 OTUs for the 660 bp and 4,854 OTUs for the 1,256 bp amplicon set (Table 1).

**Table 1.**
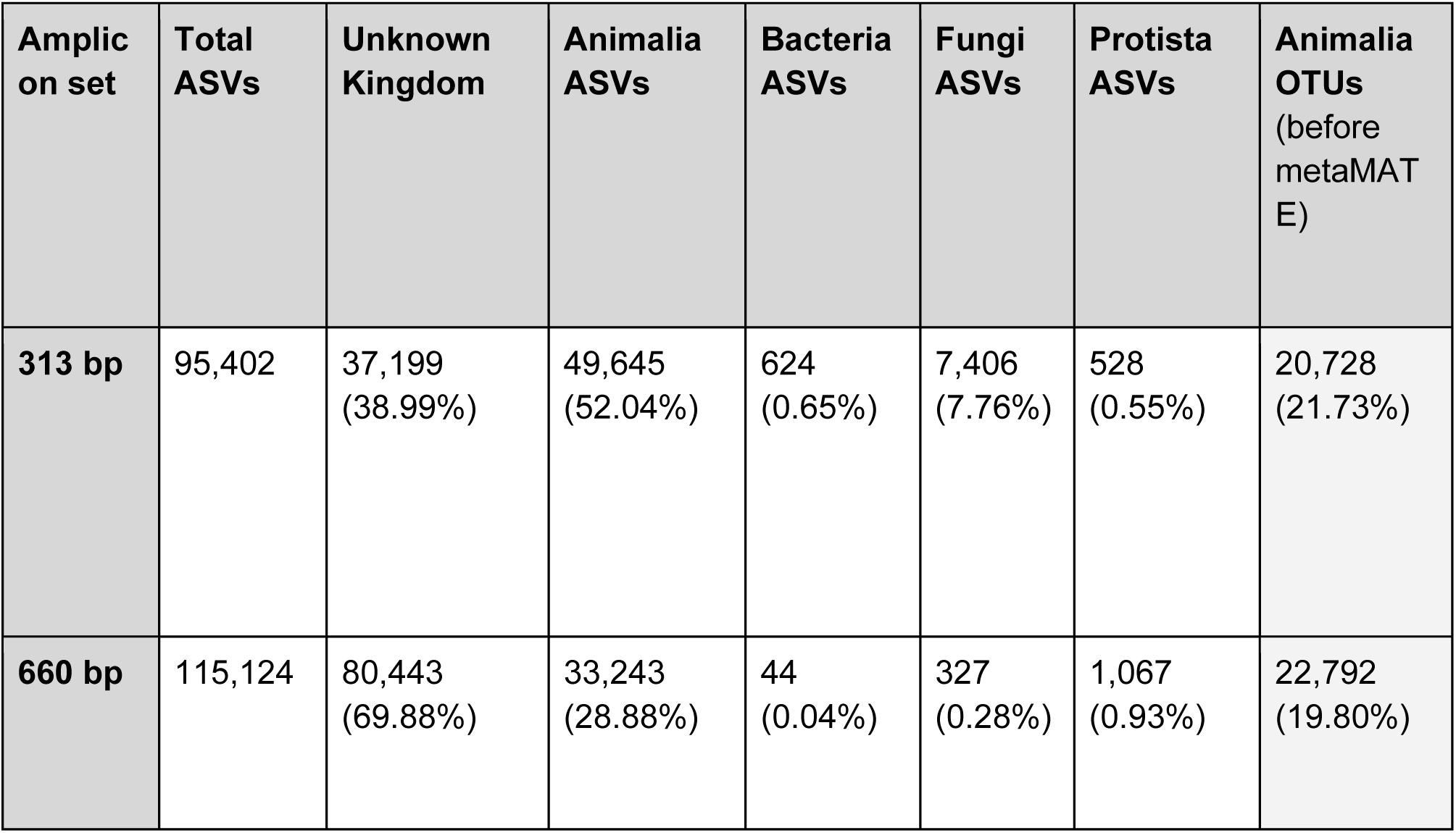
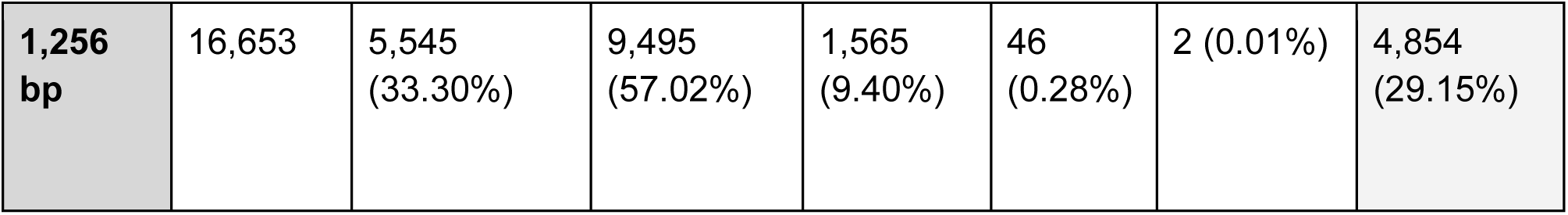
Summary of ASVs and Animalia OTUs before application of metaMATE filtering across three amplicons (313 bp, 660 bp, and 1,256 bp) from 34 soil samples. All percentages are based on the total ASV count.

### Verified authentic and non-authentic OTUs

Following metaMATE filtering, the total number of verified authentic OTUs was 434, 162 and 188 OTUs for the 313 bp, 660 bp and 1,256 bp datasets, respectively (Figure 2A). Most of the OTUs across all three datasets remained unclassified by metaMATE, ranging from 2,755 (1,256 bp amplicon) to 19,463 (313 bp amplicon) OTUs. The highest number of verified non-authentic OTUs was documented with 660 bp (n = 12,011) and the lowest with 313 bp set (n = 831). The 313 bp amplicon set also contained the lowest fraction of verified non-authentic OTUs (4%; Figure 2B).

**Figure 2.**
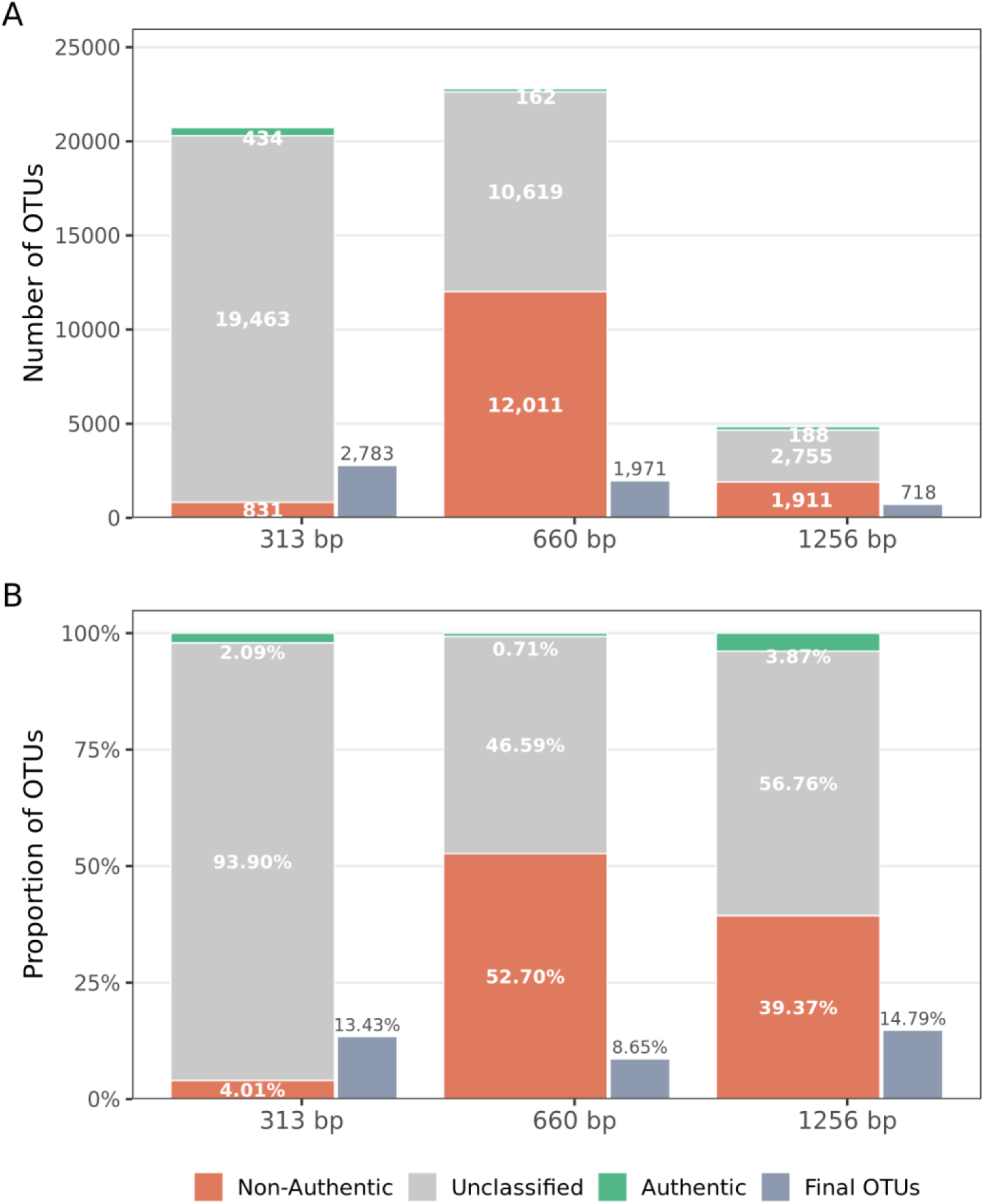
(A) Number of OTUs per amplicon set classified by metaMATE as verified authentic, unclassified and verified non-authentic, and final number of OTUs after filtering (slate bar). (B) The same data expressed as proportions of OTUs within each amplicon set.

The abundance filtering thresholds applied with metaMATE were largely variable among datasets. For the 313 bp amplicon set, the abundance thresholds applied per sample ranged between 33 to 143 reads (mean of 83) to remove at least 95% of verified non-authentic sequences (see Methods). In contrast, thresholds for 660 bp and 1,256 bp amplicon sets ranged between 1 and 13 reads (mean of 2 for 660 bp and 3 reads for 1,256 bp). Noise filtering with metaMATE reintegrates all authentic sequences (herein, OTUs) regardless of their read abundance. After the removal of verified non-authentic OTUs and reinclusion of verified authentic OTUs, 2,783 OTUs (13.4%) were retained from the initial 20,728 OTUs for the 313 bp amplicon (Figure 2). OTU retention for 660 bp and 1,256 bp amplicon sets was 8.7% (1,971 OTUs) and 14.8% (718 OTUs), respectively (Figure 2).

To compare OTU status assignments by metaMATE (authentic, non-authentic, unclassified), Animalia ASVs from the 313 bp and 660 bp amplicon sets, and separately from the 660 bp and 1256 bp amplicon sets, were clustered into “unified” OTUs. Clustering of the 313 bp and 660 bp sets produced 35,016 OTUs, including 2,878 OTUs shared between datasets. Agreement in OTU status was highest for the unclassified category (1,457, 73.8%; Figure 3). The highest level of disagreements involved unified OTUs assigned to the unclassified category in the 313 bp dataset, but as non-authentic in the 660 bp amplicon set (364, 18.4%; Figure 3). These 364 OTUs generally showed low taxonomic assignment confidence across ranks (Figure S3A); however, eight had high bootstrap support at the species level (≥ 0.95) for OTUs from 660 bp amplicon set, but were still classified as non-authentic. Similarly, 12 out of 18 OTUs classified as authentic in the 313 bp amplicon set, but as non-authentic in the 660 bp set (Figure 3) also had a high species level support (Figure S3B). In these cases, indels causing frameshifts and thus stop codon detection led to their classification as non-authentic (Table S2). Together, these 20 false-positive non-authentic 660 bp OTUs represented 19 species level features. Eleven species remained in the final 660 bp OTU table and these retained species had notably higher sequence abundance compared to the excluded conspecific ones (mean: 345.9 vs. 1.8 sequences per OTU). The excluded conspecific OTUs, represented by very low read abundance, suggest that these distinct OTUs are formed by low-frequency erroneous variants, whereas high-abundance conspecific OTUs represent more likely authentic mitochondrial sequences.

**Figure 3.**
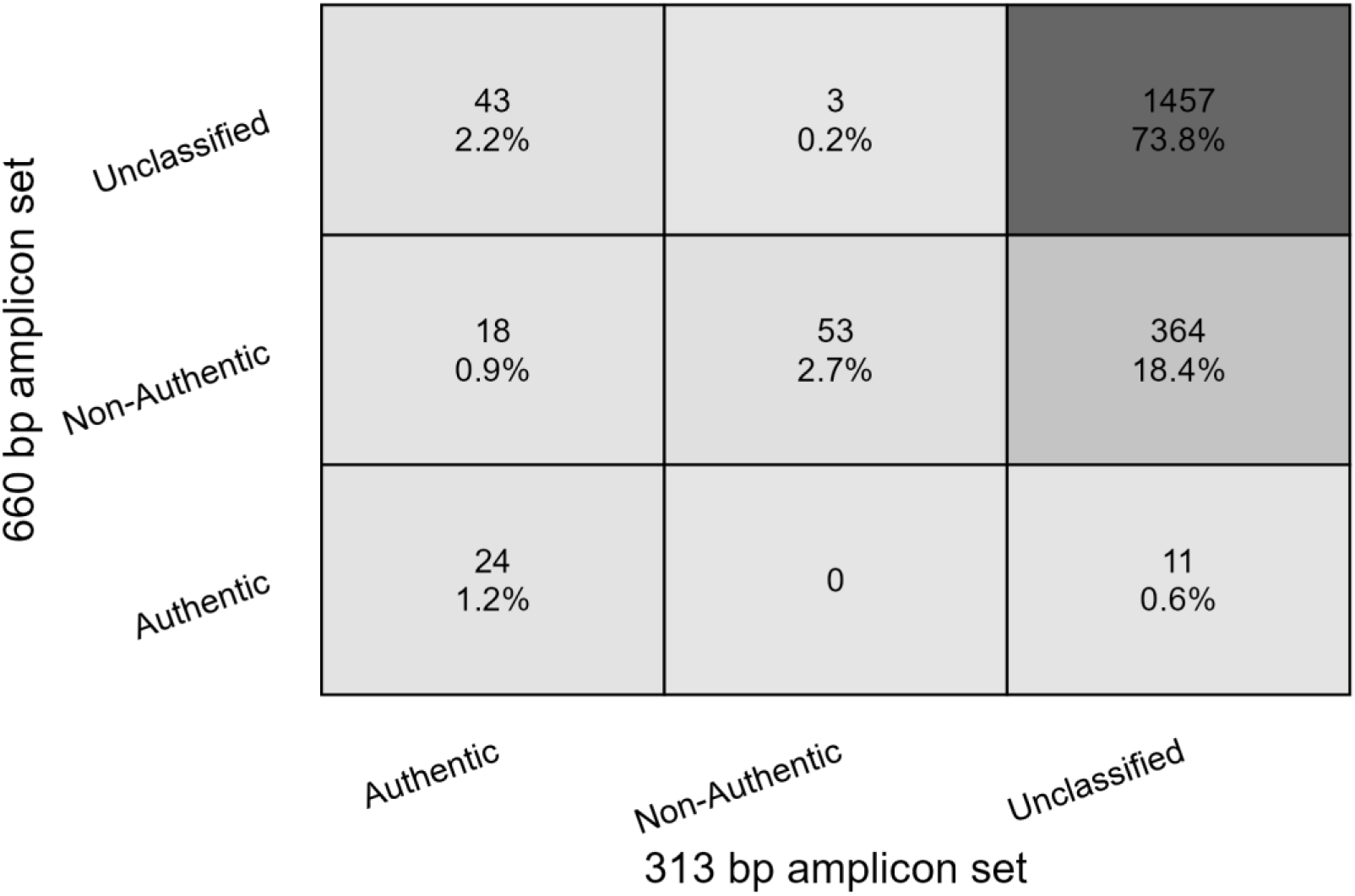
Heatmap of OTU status (assigned by metaMATE) for shared unified OTUs (n = 1,576) between 313 bp and 660 bp amplicon sets. Animalia ASVs (prior metaMATE) from 313 bp and 660 bp amplicon sets were truncated to 300 bp and clustered together to form unified OTUs.

Interestingly, 11 OTUs were classified as unclassified in the 313 bp dataset but authentic in the 660 bp dataset (Figure 3). In these cases, at least one of the ASVs in the original OTU cluster in 660 bp data showed a perfect match to the reference sequence, whereas the corresponding 313 bp ASVs sequences in the same unified cluster differed by few mismatches (thus received ‘unclassified’ status).

Although 18.4% unified OTUs (364; Figure 3) received an unclassified status with 313 bp and a non-authentic status in the 660 bp amplicon set, not all of them were retained in the final 313 bp OTU table (after metaMATE filtering). Based on abundance thresholds inferred from non-authentic OTUs in the 313 bp set, metaMATE removed 80.5% (293) of the OTUs that were suggested to be non-authentics based on the 660 bp sequences. The remaining 71 OTUs (19.5%) had a substantially higher mean sequence abundance than the removed OTUs (mean 581 vs 44 reads per OTU). Despite their higher sequence abundance, these 71 retained OTUs showed weak taxonomic support, with mean bootstrap values of 0.79 at the Phylum level and 0.13 at species level. This suggests that many OTUs that are unclassified to a Phylum level (with a cutoff of 0.8) may represent additional artefacts or off-target taxa.

### Taxonomy assignment confidence of filtered unified OTUs

Taxonomy assignment confidence within the 313 bp and 660 bp amplicon sets for only the shared unified OTUs (using the unclipped sequences) that were represented in the corresponding final OTU sets revealed highly similar patterns across taxonomic ranks (Figure 4). At higher ranks (phylum to class), a larger proportion of unified OTUs had highly confident assignments (exceeded the 0.95 threshold) in the 313 bp dataset, whereas towards lower ranks (order to genus) the 660 bp dataset showed slightly higher fraction of OTUs with bootstrap support ≥ 0.95 (Figure 4A).

**Figure 4.**
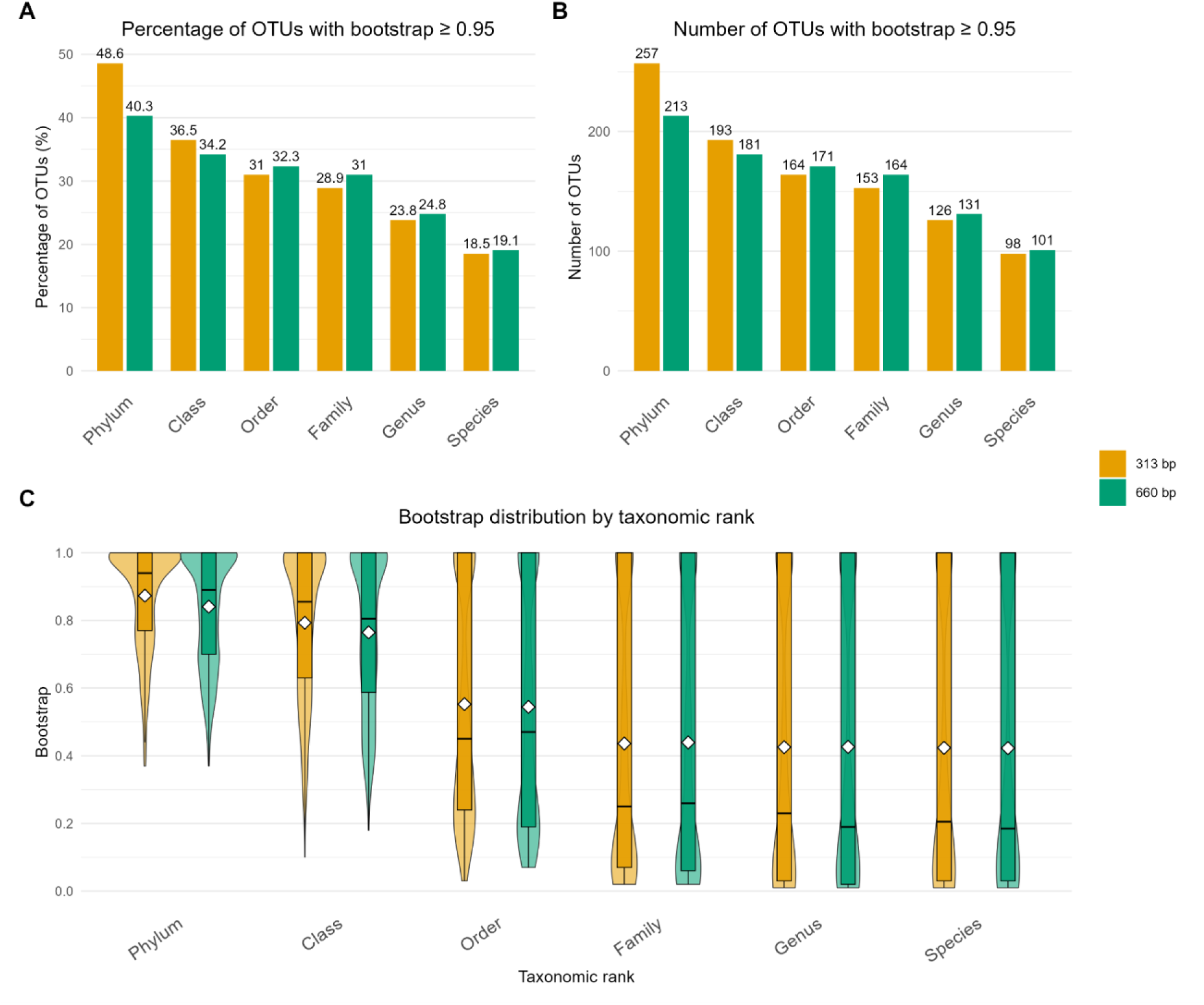
Taxonomy assignment confidence (bootstrap) comparison for shared unified OTUs (313 bp and 660 bp amplicon sets) that were present in both final OTU sets (n = 529). (A) Percentage of OTUs with SINTAX bootstrap values ≥ 0.95 at each taxonomic rank. Values on top of bars represent the percentage of OTUs meeting the threshold. (B) Absolute number of OTUs with bootstrap values ≥ 0.95 at each taxonomic rank. (C) The distribution of bootstrap values where boxplots indicate medians and interquartile ranges and diamonds mark mean bootstrap values.

Across unified OTUs that were shared between the 660 bp and 1,256 bp amplicon sets (n = 247), the 660 bp amplicons showed consistently higher bootstrap support and more high-confidence (≥0.95) assignments (Figure S4). Because the reference database is dominated by ∼650 bp COI barcode sequences (mean length 658.3 bp), many sequence patterns present in the 1,256 bp reads are absent or poorly represented in the reference database. As a result, the classifier may assign lower confidence, especially at deeper taxonomic ranks.

### False-positive and false-negative chimeras

The final OTUs (after metaMATE filtering) were analyzed with the BlasCh module to detect any remaining chimeras (false negative chimeras) in the final datasets. No multiple alignment matches (that is, chimeric sequences) were identified across all three datasets. However, comparison of the ASVs that were discarded during the chimera filtering process, against a reference database (BlasCh module analysis), revealed that the proportion of false-positive chimeras accounted for 16.5% (n = 4,810), 10.0% (n = 254), and 2.3% (n = 2) for the 313 bp, 660 bp, and 1,256 bp amplicon sets, respectively (Figure S5). False positive chimeras are ASVs with reliable matches against reference sequences (>95% identity and coverage) but incorrectly flagged as chimeric by the chimera filtering algorithm. The precision rate for chimera classifications was thus highest for 1,256 bp (0.97), followed by the 660 bp (0.90) and 313 bp (0.83). This indicates that longer sequences improve the ability to distinguish chimeric sequences from authentic ones.

### OTU richness and community composition across amplicon sets

Although the total OTU richness was highest in the 313 bp set final OTU table (Figure 2A), pairwise comparisons of per-sample OTU richness revealed no significant difference between the 313 bp and 660 bp amplicon sets (Wilcoxon rank-sum test with Bonferroni correction: p = 0.389; Figure 5A). The 1,256 bp amplicon set exhibited significantly lower OTU richness compared with both shorter amplicon sets (p < 0.001; Figure 5A). The 660 bp amplicon set yielded the highest mean OTU richness per sample (mean ± SD: 185 ± 29.1 OTUs, range 139-241), followed by the 313 bp amplicon (174 ± 52.9 OTUs, range 86-320), and the 1,256 bp amplicon (65.6 ± 22.6 OTUs, range 40-122; Figure 5A). Compared with the 660 bp dataset, the 313 bp dataset contained substantially more OTUs occurring only in one sample: 1,709 (61.4% of all 313 bp OTUs) vs. 1,080 (54.8% of all 660 bp OTUs). This indicates that the higher total richness observed in the 313 bp dataset (Figure 2A) was mainly driven by OTUs that occurred in only a single sample, which contributed substantially to total richness but comparatively little to average per-sample richness.

**Figure 5.**
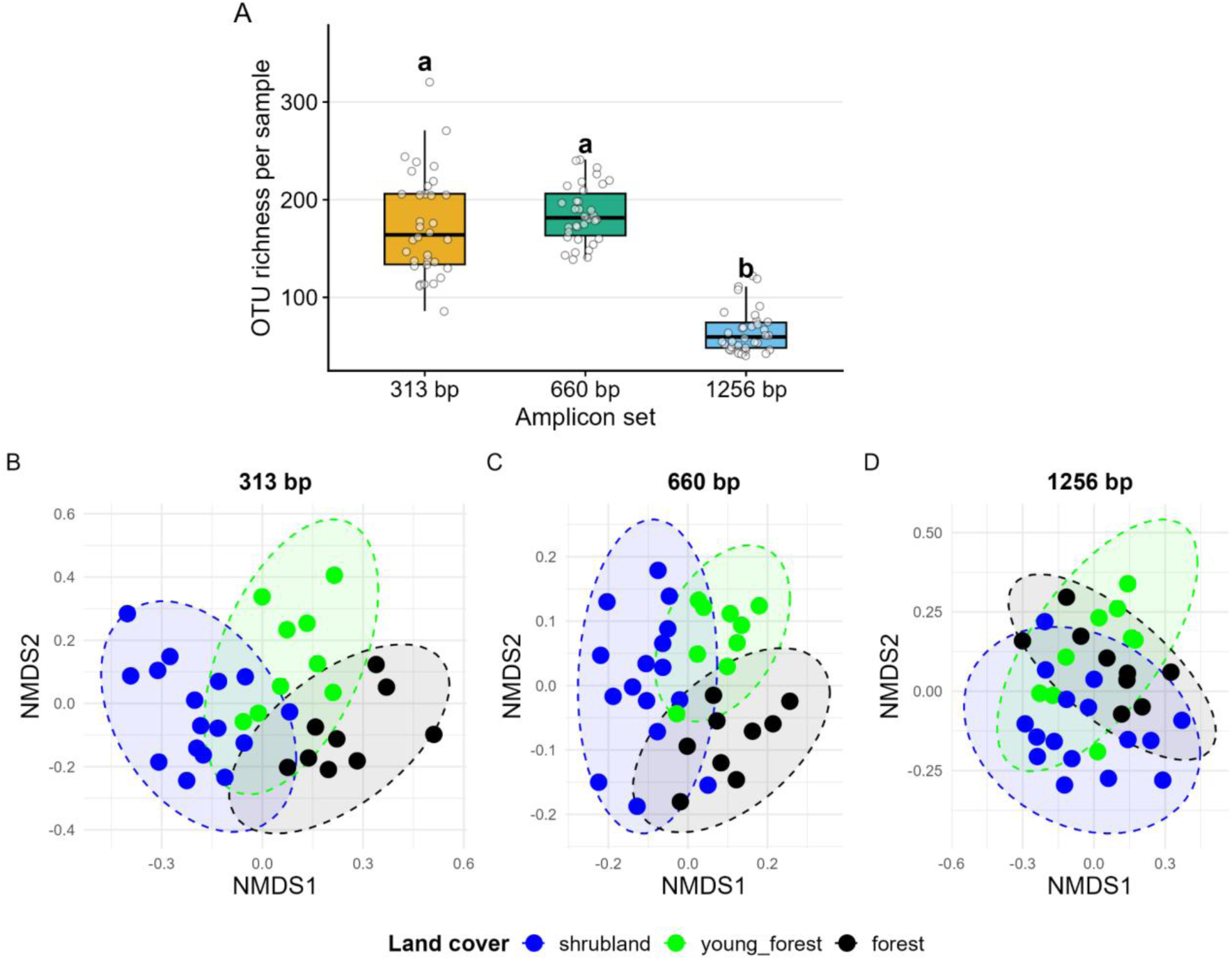
A) OTU richness per sample across the three COI amplicon sets (313 bp, 660 bp, 1,256 bp; n = 34 per amplicon set). Boxplots show the median (centre line), interquartile range (box), and 1.5 × interquartile range (whiskers) of the number of OTUs per amplicon set. Different letters above boxplots indicate significant pairwise differences between amplicon sets in OTU richness per sample. B-D). Ordination plots (NMDS) of soil invertebrate community composition based on Jaccard dissimilarities (on presence-absence data) for the three COI amplicon sets: 313 bp (B), 660 bp (C), and 1,256 bp (D); each point represents one soil sample (n = 34), coloured by land cover type. Dashed ellipses represent 95% confidence ellipses for each land cover group.

Procrustes’ analysis revealed significant concordance between all three COI amplicon sets in their recovery of community compositional patterns (Figure 5B-D). The strongest agreement was observed between the ordinations from 313 bp and 660 bp sets (Procrustes R = 0.637, p < 0.001), followed by the two PacBio datasets (660 bp vs. 1,256 bp; Procrustes R = 0.590, p < 0.001) and the 313 bp and 1,256 bp amplicon sets (Procrustes R = 0.371, p < 0.001). Consistent with these patterns, taxonomic composition at the order level was most similar between the 313 bp and 660 bp amplicon sets, whereas the 1,256 bp dataset showed the greatest divergence in relative composition (Figure S6).

## Discussion

In this study, we examined the comparative performance of a commonly used COI mini-barcode (313 bp), the standard full-length animal COI barcode (658 bp, here represented with 660 bp), and a longer COI amplicon (1,256 bp) for soil eDNA metabarcoding. The longer amplicon sets, 660 bp and 1,256 bp, yielded the most effective detection of non-authentic OTUs and filtering in terms of required additional abundance thresholds to remove most of the estimated noise. Additionally, the chimera filtering process was more accurate with the longer sequences by filtering out less false-positive chimeras compared with short (313 bp) sequences. While the 1,256 bp amplicon represents a longer sequence of the COI region, its benefits were constrained in practice, mostly because reference sequences rarely extend beyond the standard barcoding region (>658 bp). Although the mini-barcode (313 bp) and standard barcode (660 bp) COI regions yielded largely comparable results in terms of community composition, the 660 bp amplicons facilitated more effective filtering of spurious sequences, thereby representing an optimal target region for eDNA metabarcoding.

### OTU authenticity across amplicon sets

In our study, the shortest amplicon (313 bp) had the highest number of retained OTUs, as well as the highest number of verified authentic OTUs. At first glance, this could suggest that mitochondrial templates were amplified more efficiently for the shorter fragment. However, when compared with the longer amplicons, this pattern raises an important question: does the higher number of OTUs reflect genuine success or instead result from a reduced ability to identify non-authentic sequences at this fragment length? Because shorter sequences contain fewer informative sites, the probability of detecting mutations indicative of a non-authentic sequence is inherently lower (Schultz & Hebert 2022, Porter & Hajibabaei 2021; Hebert et al. 2023; Edgar et al. 2011). As a result, artefactual sequences lacking obvious diagnostics within the 313 bp region may be more likely to pass filtering steps. Direct comparison of OTU status assignments between the 313 bp and 660 bp datasets supports this interpretation. Among shared OTUs, the most frequent disagreement involved sequences that were classified as non-authentic in the 660 bp set, but remained unclassified in the 313 bp set. A large majority of these OTUs exhibited low taxonomic assignment confidence (Figure S3), further suggesting that they may rather represent artefactual sequences, and that their unclassified status in the shorter amplicon set is reflective of limited diagnostic power. Nevertheless, metaMATE filtering demonstrated to be effective in estimating the remaining noise in the 313 bp amplicon set and removed the majority of the sequences flagged as non-authentic with longer reads.

Although longer amplicons are generally more effective for NUMT detection (Porter & Hajibabaei 2021; Schultz & Hebert 2022), they may also be more susceptible to indel-associated sequencing errors – a known limitation of PacBio sequencing (Wenger et al. 2019). Indels can introduce frameshifts that result in premature stop codons or otherwise nonsensical translations during open reading frame analysis. Accordingly, we identified a few instances in which OTUs from the 660 bp dataset were wrongly classified as NUMTs because of indel induced frameshifts. The occurrence of detected false positive non-authentic OTUs was low, suggesting that the impact of such errors on downstream analyses is minimal. However, when operating at the ASV level, the impact of false-positive non-authentic ASVs (as well as indel-induced artefactual ASVs) may be higher.

The longest fragment (1,256 bp) yielded fewer (including proportionally) non-authentic OTUs than the 660 bp set, but had a proportionally higher rate of unclassified OTUs. This suggests that targeting longer sequences may preferentially amplify valid mitochondrial templates that remain unclassified (due to the missing reference sequences) while excluding nuclear copies that are shorter (e.g. Marshall and Parson 2021, Tao et al. 2023). Additionally, longer amplicons also improved the detection of other artefacts, as for example, the higher precision of chimera classification indicates that increased sequence length provides more reliable signals for distinguishing recombination artefacts from genuine biological sequences (Edgar et al. 2011; Figure S5).

### COI amplicon length implications for taxonomic resolution

One motivation for targeting longer amplicons is their ability to improve taxonomic assignment (Doorenspleet *et al*. 2025; Tedersoo *et al*. 2021). Assuming that reference databases contain representative sequences for the taxa present in the query dataset, longer query sequences should increase reliability of taxonomic assignments (Clarke et al. 2017, Porter & Hajibabaei 2018).

However, among the shared OTUs, the 660 bp amplicon set exhibited only a marginal increase in assignment confidence at lower taxonomic ranks while the overall confidence distributions remained highly similar to those of the 313 bp amplicon. This aligns with the findings of Yeo et al. (2020), who demonstrated that mini-barcodes can be just as effective for species identification as full-length COI barcodes. Therefore, in general, the 313 bp fragment captures sufficient phylogenetically informative variation, such that the additional nucleotide positions provided by the 660 bp sequence add little additional information for species identification purposes (Meusnier et al. 2008). Furthermore, when classifying longer sequences that extend beyond the length coverage of the reference database, we observed a substantial decline in assignment confidence. This pattern reflects the mismatch between query length and reference database composition. Because the reference database is largely composed of ∼650 bp COI barcode sequences, longer reads contain additional k-mers that are absent or underrepresented in the database. As a consequence, classifier consistency decreases, leading to reduced confidence in taxonomic assignments. Here, we used a k-mer based classifier (SINTAX), but a similar decline in assignment confidence is expected with alignment-based approaches such as BLAST if the overlap between query and reference sequence is included into the reliability calculation of a taxonomic assignment. Overall, these results provide further support that increasing COI amplicon length does not inherently enhance taxonomic assignment confidence. Shorter COI fragments can be sufficient for reliable identification, whereas sequences extending beyond the standard barcoding region may even reduce assignment confidence.

### OTU richness across amplicon sets

Here, we tested newly designed primers to amplify 1,256 bp of the COI region. This amplicon set demonstrated the lowest number of Animalia OTUs as well as the lowest proportion of target sequences among the three datasets (Figure 2, Figure S2). This likely reflects a substantial primer bias as the primers for generating 1,256 bp amplicons were designed based on Collembola and Acari sequences, but reduced amplification success of longer and potentially fragmented soil DNA templates may also have contributed. Despite that, the primer also captured substantial amounts of non-target DNA. Thus, despite the potential advantages of longer sequences for artefact detection, this primer pair appears poorly suited for broad soil animal eDNA metabarcoding without further optimization.

Although the 313 bp amplicon set in our study had, on average, more than 20x higher raw sequencing depth per sample than the 660 bp dataset (Figure S2, S5), the total observed OTU richness was comparable between both datasets (Figure 2), with no statistically supported differences in per-sample OTU richness (Figure 5A). This is also concurrent with the findings by Varusk et al. (2025), who reported non-significant difference in soil mite OTU richness between Illumina and PacBio datasets despite the latter having largely lower sequencing depth. This indicates that lower sequencing depth does not necessarily translate into lower richness recovery. However, this does not reflect a sequencing platform effect, but more likely the use of different primer pairs to amplify short and long reads. Here, we found that only 18.5% of 313 bp sequences (relative to the raw average sequencing depth per sample) were retained after discarding the non-Animalia ASVs (Figure S2). Similarly, previous studies have shown that the primers targeting the COI mini-barcodes recover a substantial proportion of non-metazoa taxa from eDNA samples (Bakker et al. 2019, Collins et al. 2019, Anslan et al. 2021, Couton et al. 2023). However, in our study, Animalia sequences accounted for 74.8% of reads in the 660 bp dataset, indicating that the standard Folmer primers provided more efficient enrichment of the target amplicons from eDNA. This suggests that the raw sequencing depth per sample may be naturally lower when generating a metabarcoding dataset with primers that capture the target group more efficiently. Therefore, the concern of sequencing cost for long-reads may be alleviated by using a lower sequencing depth per sample without compromising the recovery of target taxa.

## Conclusion

As sequencing technologies move beyond the read-length constraints that originally favoured mini-barcodes, our results argue for a transition to the standard COI barcode (∼658 bp) as an optimal marker for metabarcoding soil animal communities from eDNA. It provides an optimal balance between target sequence recovery, effective NUMT and chimera detection, and compatibility with existing reference databases for reliable taxonomic assignment. Extending beyond the standard barcode region currently offers limited additional benefit for metabarcoding applications. Mini-barcodes typically yield higher OTU richness, but also higher proportions of non-Animalia sequences, and processing of short reads may be more prone to retaining artefactual sequences. We note that recovery of the standard barcode region with PacBio sequencing may be subject to indel-related errors, albeit to a limited extent. We suggest that the application of recently developed 2 x 500 bp sequencing with the Illumina platform to sequence the full-length standard COI region would likely adequately address this error, but this is yet to be evaluated.

## Acknowledgements

This work was supported by the Research Council of Finland (Decision number 362828), Researchers Supporting Project (HCPNU2026R402) at Princess Nourah bint Abdulrahman University (Riyadh, Saudi Arabia), by the European Regional Development Fund and the program Mobilitas Pluss (MOBTP198), and by Ministry of Education and Research of Estonia (Center of Excellence AgroCropFuture, TK200). This work has been carried out within the framework of the doctorate in bioinformatics of the Universitat Autònoma de Barcelona (UAB). The authors acknowledge CSC–IT Center for Science, Finland, for computational resources, Novogene and Norwegian Sequencing Centre (NSC) for sequencing services.

## Ethics and integrity policies

Soil samples analyzed in this study are a subset of those collected by Anslan et al. (2021); sampling was conducted in accordance with relevant local and national legislation, and no specific permits were required for collection of soil for environmental DNA analysis.

## Data availability

The raw sequencing data is available in the Sequence Read Archive: PRJNA743174 (used BioSample names in Table S1).

## Author Contributions

Conception of the work: SA, MHE

Acquiring of funds: SA, KMA

Designed new analytical tools: MHE, CA, PA, BCE, SA

Data generation: SA, SV, KS, LT, KMA

Data analysis: MHE, SA, AH, MM

First draft of the manuscript: SA, MHE

Review and edition of manuscript: MHE, SA, SV, KS, AH, MM, LT, KMA, PA, CA, BCE

