## Supporting Information for "Full-length COI barcodes improve eDNA metabarcoding data denoising relative to mini-barcodes"

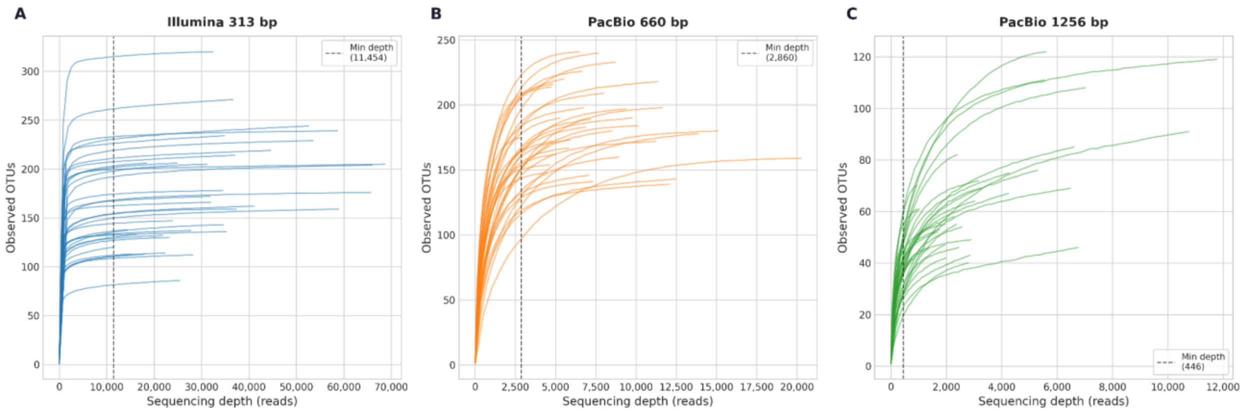


**Figure S1**. Rarefaction curves for OTU richness across the three COI amplicon datasets (A) 313 bp, (B) 660 bp (C) and 1,256 bp. Curves show the relationship between sequencing depth (x-axis) and observed OTU richness per sample (y-axis), based on the final OTU tables after metaMATE filtering. Each line represents an individual sample. Curves were generated using the scikit-bio package and visualized in Python with matplotlib.

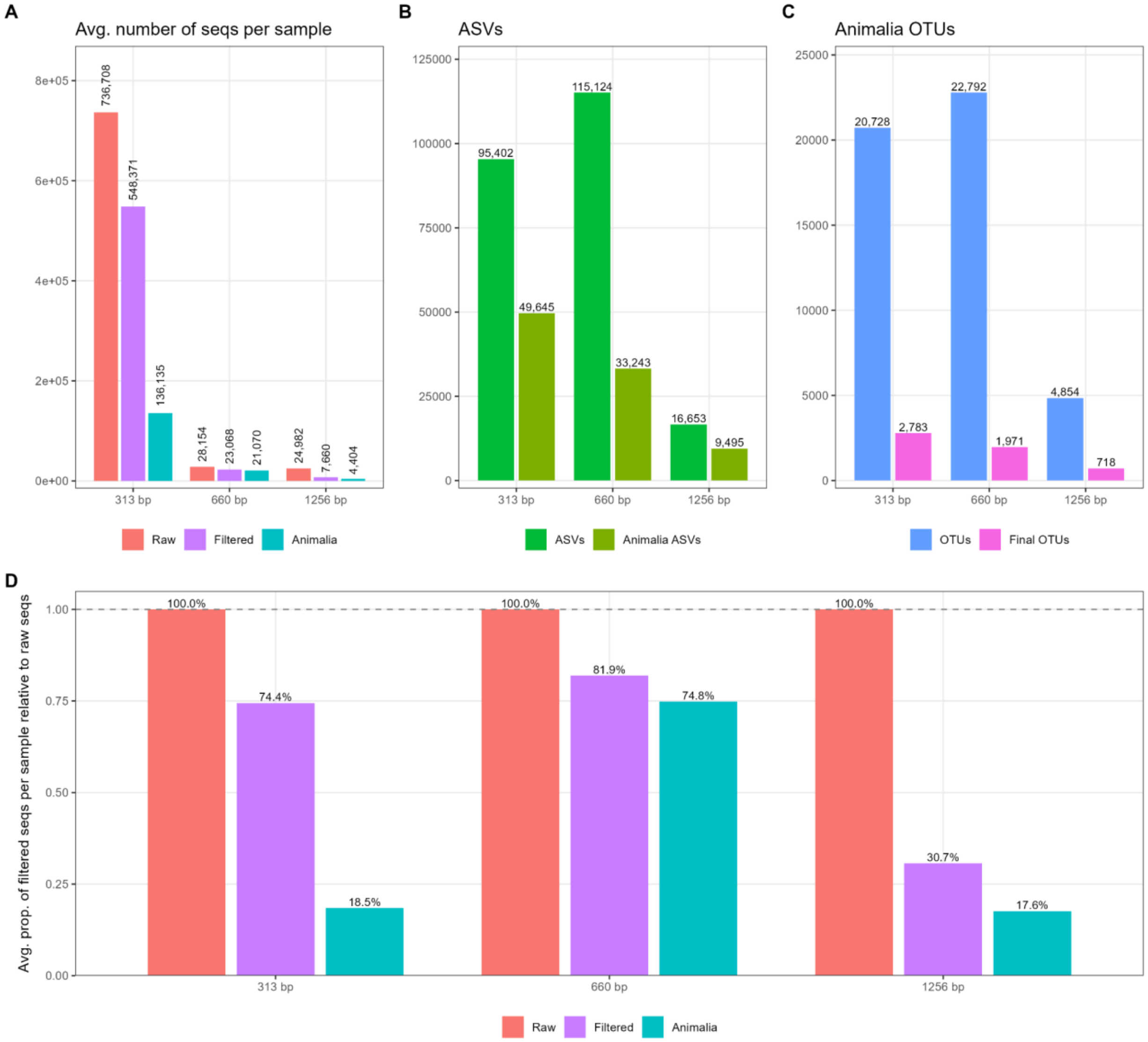


**Figure S2**. Summary of sequence processing outcomes across amplicon sets. The top left panel (Seq depth) shows bar plots for average sequence depth per sample for raw (red bars) and DADA2 filtered data (cyan bars). The bottom panel shows the average proportion of sequences retained after DADA2 filtering (cyan bars) relative to raw sequencing depth (red bars); numbers above bars indicate the corresponding sequence counts. The top middle panel (ASVs) shows ASV counts per amplicon set after the DADA2 pipeline (and before metaMATE; green bars). The top right panel shows OTU counts before (light blue bars) and after metaMATE filtering (magenta colored bars).

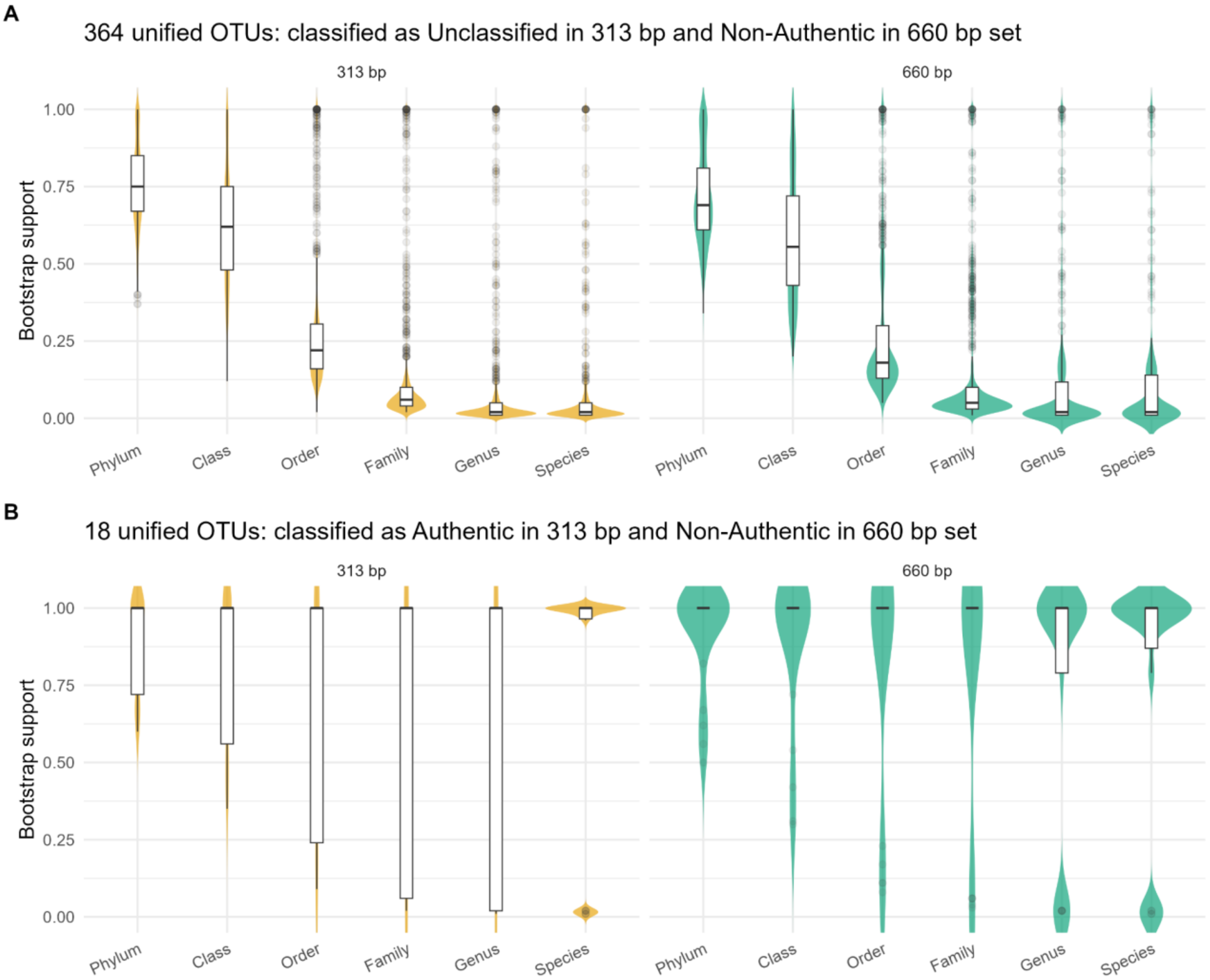


**Figure S3**. Distribution of taxonomy bootstrap support values for ASVs belonging (A) to unified OTUs classified as unclassified in the 313 bp dataset and non-authentic in the 660 bp dataset (n = 364 unified OTUs); and (B) to unified OTUs classified as authentic in the 313 bp dataset but non-authentic in the 660 bp dataset (n = 18 unified OTUs). Violin plots show the full distribution of bootstrap support at each rank for each amplicon set (panels 313 bp and 660 bp). Overlaid boxplots display the median and interquartile range; points represent individual ASV bootstrap values.

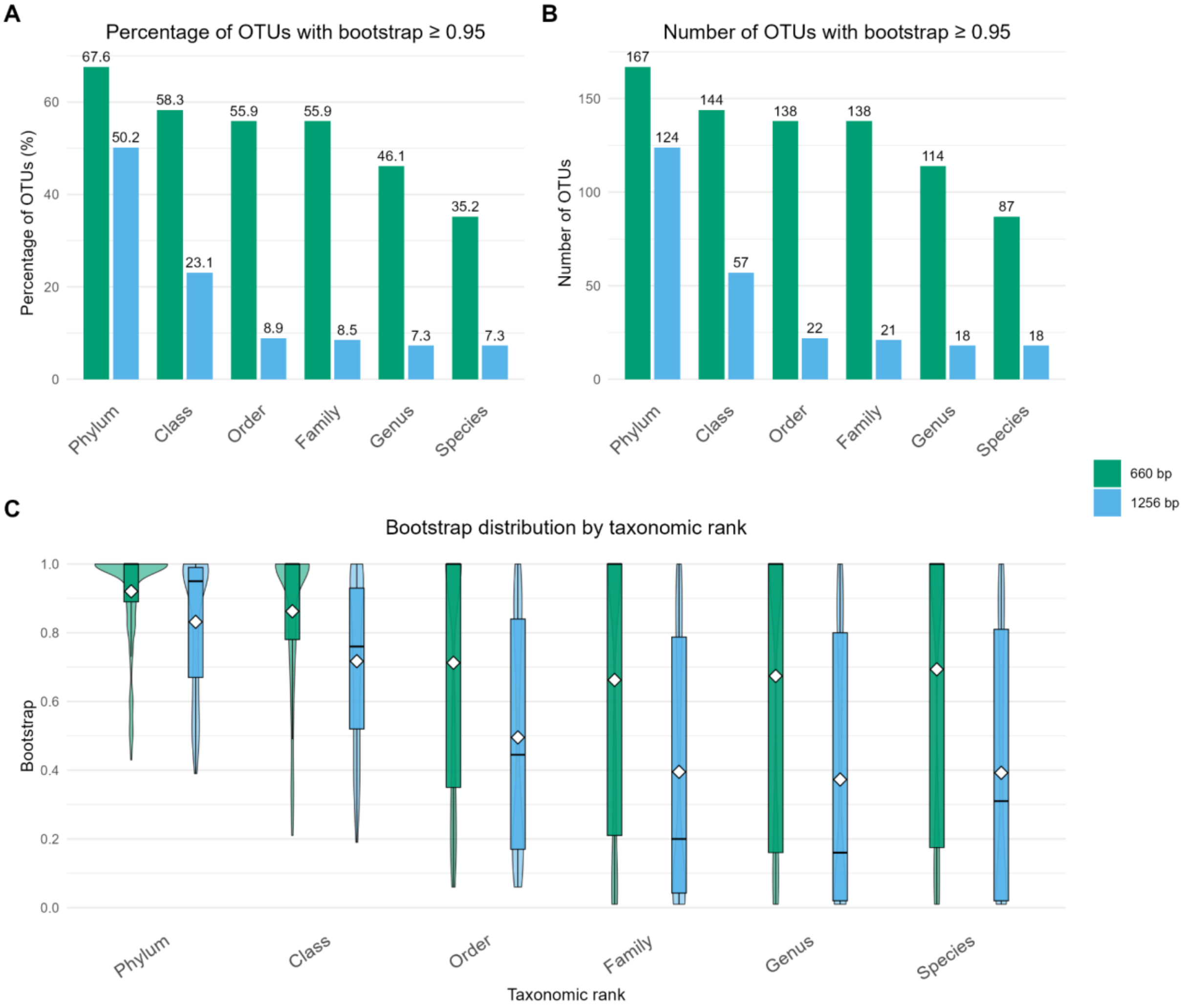


**Figure S4**. Distribution of taxonomy bootstrap support values for ASVs belonging to unified OTUs shared between the 660 bp and 1,256 bp datasets (n = 247 unified OTUs that are also represented in the final OTU tables). Violin plots show the full distribution of bootstrap support at each taxonomic rank for each amplicon set (panels 660 bp and 1,256 bp). Overlaid boxplots display the median and interquartile range; points represent individual ASV bootstrap values. Clustering of 660 bp and 1,256 bp ASVs (with uniform sequence length, see Methods) formed a total of 26,329 OTUs with 2,878 OTUs formed by ASVs from both datasets, and 247 unified OTUs were represented in both 660 bp and 1256 final OTU tables.

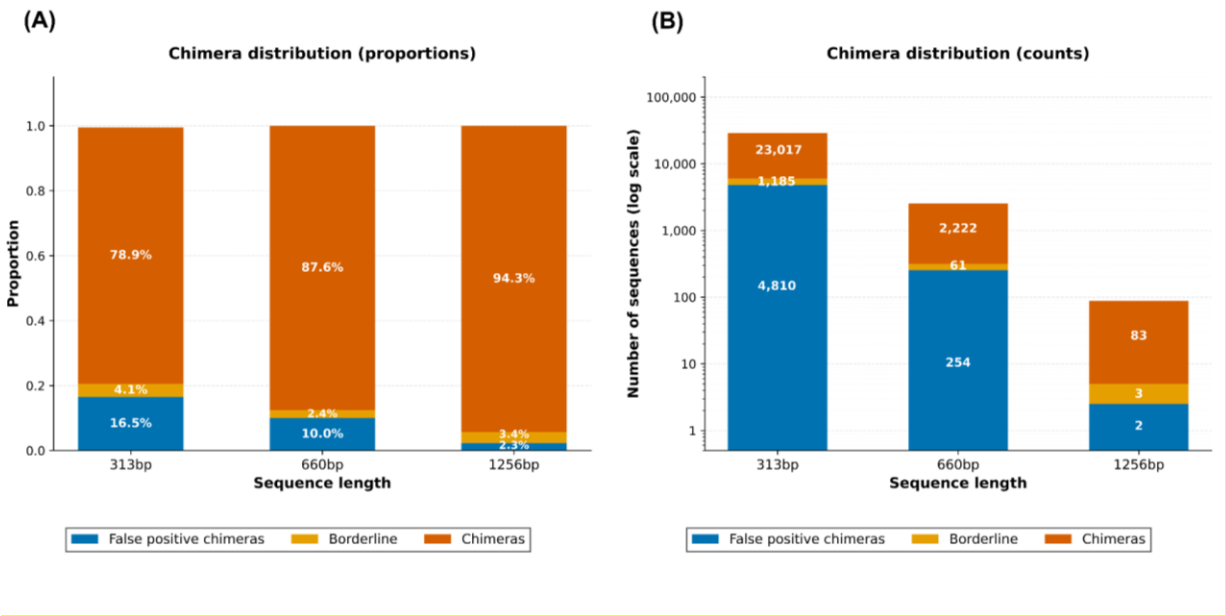


**Figure S5.** Chimera distribution across COI amplicon sets of different lengths based on BlasCh results. (A) Proportional composition of discarded sequences classified as true chimeras, borderline sequences, and false-positive chimeras for the 313 bp, 660 bp, and 1,256 bp amplicon sets. (B) Absolute counts (log scale) of the same sequence categories across amplicon sets. Query ASVs that demonstrated at least 95% sequence identity with at least 95% query coverage against reference sequences were considered as false-positive chimeras. ASVs with at least 80% identity and 89% query coverage against a reference database sequence were considered “borderline” chimeric ASVs.

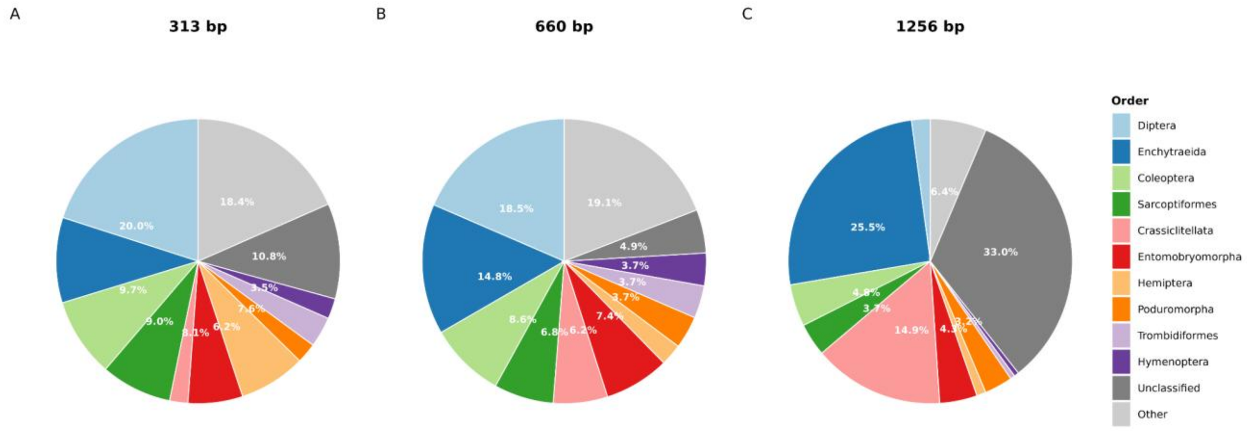


**Figure S6.** Distribution of authentic OTUs across taxonomic orders for three amplicon lengths (A) 313 bp, (B) 660 bp, and (C) 1,256 bp. Taxonomic assignments were retained only when bootstrap support was $\geq 0.8$; OTUs not meeting this threshold were categorized as unclassified.
